# Peptide Arrays of Three Collections of Human Sera from Patients Infected with Mosquito-Borne Viruses

**DOI:** 10.1101/650796

**Authors:** Maria del Pilar Martinez Viedma, Nurgun Kose, Leda Parham, Angel Balmaseda, Guillermina Kuan, Ivette Lorenzana, Eva Harris, James E. Crowe, Brett E. Pickett

## Abstract

Global outbreaks and epidemics caused by emerging or re-emerging mosquito-borne viruses are becoming more common. These viruses belong to multiple genera including *Flavivirus* and *Alphavirus* and often cause non-specific or asymptomatic infection, which can confound viral prevalence studies. In addition, many acute phase diagnostic tests rely on the detection of viral components such as RNA or antigen. Standard serological tests are often not reliable for diagnosis after the acute phase due to cross-reactivity among viruses (*e.g*. flaviviruses). In order to contribute to development efforts for mosquito-borne serodiagnostics, we incubated 137 human sera on individual custom peptide arrays that consisted of over 866 unique peptides in quadruplicate. Our bioinformatics workflow to analyze these data incorporated machine learning, statistics, and B-cell epitope prediction. The unprocessed array data can be useful in separate meta-analyses that can be applicable to diverse efforts including the development of new pan-flavivirus antibodies, more accurate epitope mapping, and vaccine development against these viral pathogens.

## INTRODUCTION

Zika virus (ZIKV) is an arbovirus that belongs to the *Flavivirus* genus within the *Flaviviridae* family. In addition to ZIKV, many other mosquito-borne viruses exist that negatively affect public health, including dengue (DENV) and chikungunya (CHIKV) viruses, among others. ZIKV is primarily transmitted by the bite of infected *Aedes* spp. mosquitoes, with limited instances of sexual transmission also being reported [1-4]. The recent worldwide epidemic has demonstrated that ZIKV is a neuropathic virus that is associated with fetal microcephaly and other congenital defects in infected pregnant women, and Guillain-Barré Syndrome in adults [5]. Due to the number of ZIKV infections in recent years and the continued threat of ZIKV re-emerging around the world, there is still an urgent need for rapid and accurate surveillance assays in order to rapidly identify new outbreaks. Distinguishing between infection with multiple co-circulating arboviruses that have similar clinical signs and symptoms makes accurate prevalence calculations and diagnosis extremely difficult—especially after convalescence [6-10].

The sequence similarity among many of these flaviviruses at the protein level contributes to the observed cross-reactivity in serological assays, which is especially high in the E protein and also present in the NS1 protein [11]. Although reports showing antibodies against other viral proteins are detectable [12], the E and NS1 proteins are the primary targets of the humoral anti-flavivirus immune response in humans [13-15].

Recent efforts to generate whole-genome sequences for these pathogens enable the application of bioinformatics tools to mine the data for trends and patterns that can be clinically applicable [16-20]. The meta-CATS (metadata-driven Comparative Analysis Tool for Sequences) algorithm is a statistical workflow that rapidly identifies sequence variations that significantly correlate with the associated metadata for two or more groups of sequences [21]. This algorithm has been used previously to identify residues within 15-mer linear peptide regions that have high predicted specificity and sensitivity values and that could therefore be useful for detecting antibodies against a variety of mosquito-borne *Flavivirus* species [22]. Quantifying the reactivity of this set of peptides using high-throughput custom peptide arrays enabled the efficient and simultaneous testing of all peptides against each serum sample with higher efficiency than what is possible with manual ELISA-based technology [23].

The data presented here are the product of combining upstream computational methods to predict peptides capable of distinguishing each virus with downstream high-throughput screening of relevant sera using peptide arrays. We have recently completed an analysis of 137 serum samples using peptide arrays (each containing 866 experimental viral peptides) to identify 15-mer linear peptides that could be useful as serodiagnostic reagents to detect prior infection with mosquito-borne viruses. Specifically, we tested peptides representing different co-circulating mosquito-borne viruses including: Zika, dengue 1-3, West Nile and chikungunya viruses. Applying machine learning, a weighting scheme, and B-cell epitope prediction algorithms to these data enabled us to identify pools of 8-10 peptides that are predicted to be immunodominant across human sera from individuals infected in Central and South America. In addition, we have separately evaluated these peptides using a set of well-characterized sera. These data could be used by the scientific community to develop improved serological diagnostic methods for detecting past infection with one or more of these viral pathogens.

## METHODS

### Peptide preparation and microarray printing

A subset of the previously-predicted diagnostic peptides [21], representing multiple mosquito-borne virus species and subtypes, were synthesized at the Center for Protein and Nucleic Acid Research at The Scripps Research Institute (TSRI) [23, 24]. This selected collection of peptides consisted of 15-mers with sequences that represented the consensus amino acid sequence among strains belonging to each of our six target taxa including: CHIKV, DENV1, DENV2, DENV3, WNV, and ZIKV. Peptides on the array that represented mosquito-borne virus taxa for which there were no serum samples were ignored in downstream quantification and computation. As such, a total of 25, 51, 28, 34, or 70 peptides in the E protein as well as 15, 19, 15, 23, or 70 peptides in the NS1 protein (all derived from DENV1, DENV2, DENV3, WNV, or ZIKV sequences, respectively) were evaluated in these experiments. A set of 25 peptides spanning portions of the CHIKV E2 protein that had previously been reported as relevant for detecting anti-CHIKV antibodies were also included [25]. Synthesized peptides were suspended in 12.5 μL DMSO and 12.5 μL of ultra-pure water. Immediately prior to printing, suspended peptides were diluted 1:4 in a custom protein printing buffer [Saline Sodium Citrate (SSC) 300 mM sodium citrate, pH 8.0, containing 1 M sodium chloride and supplemented with 0.1% Polyvinyl Alcohol (PVA) and 0.05% Tween 20], in a 384-well non-binding polystyrene assay plate. Two positive control peptides, hemagglutinin A (HA) (YPYDVPDYA) and FLAG tag (DYKDDDDK), together with a dye that permanently fluoresces at 488 nm (Alexa Fluor 488) were included in the print to guide proper grid placement and peptide alignment, as well as to serve as printing controls as well as controls to quantify the maximum fluorescence for the assays.

Quadruplicate sets of all peptides were printed onto N-hydroxysuccinimide ester (NHS-ester) coated NEXTERION Slide H (Applied Microarrays) slides at an approximate density of 1 ng/spot, using a Microgrid II (DigilabGlobal) microarray printing robot equipped with solid steel (SMP4, TeleChem) microarray pins. Humidity was maintained at 50% during the printing process. Immediately prior to interrogating the arrays, slides were blocked for 1 h with ethanolamine buffer to quench any unreacted NHS-ester on the slide. All slides were used within 2 months of printing and were stored at −20°C [23].

### Serum sources

Spent diagnostic serum samples were provided by collaborators working under three separate studies in Honduras, the United States, and Nicaragua. These sera were collected from a total of 137 consented human patients under IRB supervision and were characterized as positive for antibodies against at least one of: ZIKV, DENV1, DENV2, DENV3, WNV, and/or CHIKV.

Thirty-two deidentified plasma samples from patients suspected of Zika, chikungunya or dengue in Honduras were obtained at the discretion of health care providers at the Hospital Escuela Universitario from patients (ages 6-73 years old). These samples were sent to the Centro de Investigaciones Geneticas at the Universidad Nacional Autonoma de Honduras in Tegucigalpa, Honduras for ZIKV, CHIKV and/or DENV molecular testing. Twenty-three of these patients had infection with DENV and nine had infection with ZIKV confirmed by RT-qPCR during the acute phase. Convalescent samples were collected from these patients 10-30 days post-onset of symptoms between June 1 to November 30, 2016 and were tested on the custom arrays.

Seventy-three de-identified human serum samples were obtained from the Vanderbilt Vaccine Center Biorepository. Sera from individuals with previous history of natural infection with DENV, WNV, CHIKV, or ZIKV (confirmed by serology or RT-qPCR) while traveling in the Caribbean, Central or South America, or West Africa were included on arrays. For WNV, sera were from individuals with confirmed previous history of natural infection contracted during an outbreak in 2012 in Dallas, TX. The samples were collected in the convalescent phase, months to years after post-onset of symptoms.

Thirty-two de-identified human sera were collected from the Pediatric Dengue Cohort Study in Managua, Nicaragua [26, 27]. Early convalescent-phase samples were collected 15-17 days post-onset of symptoms from 9 Zika cases that were confirmed as positive for ZIKV infection by real-time RT-qPCR between January and July, 2016. Late convalescent samples were obtained from 21 DENV-positive cohort participants after RT-qPCR confirmed DENV1 (n=7), DENV2 (n=8), or DENV3 (n=6) infection and 2 DENV-negative subjects, all in 2004-2011, prior to the introduction of ZIKV to Nicaragua. Samples were analyzed by Inhibition ELISA [28, 29] and neutralization assay [30, 31]. The PDCS was approved by the IRBs of the University of California, Berkeley, and Nicaraguan Ministry of Health. Parents or legal guardians of all subjects provided written informed consent; subjects 6 years old and older provided assent.

### High-throughput screening and quantification of characterized patient sera

The 137 characterized sera were separately subjected to high-throughput screening using the synthesized peptide arrays. Sera were tested for IgG reactivity using the custom peptide array at TSRI. For immunolabeling, the incubation area around the printed grids was circumscribed using a Peroxidase Anti-Peroxidase (PAP) hydrophobic marker pen (Research Products International Corp) and the subsequent steps were performed in a humidified chamber at room temperature on a rotator. Control anti-HA and anti-FLAG monoclonal antibodies were assayed at a concentration of 10 μg/ml while 10 microliters of human sera were diluted 1:200 in PBS buffer containing Tween (PBS-T) and incubated for 1 h followed by three washes in PBS buffer. The arrays were then incubated for 1 h with goat anti-human IgG tagged with Alexa Fluor® 488 (Invitrogen) as secondary antibody. Arrays were washed three times in PBS-T, two times in PBS, and another two times in deionized water and centrifuged to dry at 200 × g for 5 mins.

The fluorescence of the processed slides was quantified using a ProScanArray HT (Perkin Elmer) microarray scanner at 488nm and 600 nm, and images were saved as high-resolution TIF files. The Imagene® 6.1 microarray analysis software (BioDiscovery) calculated the fluorescence intensity of the area within the printed diameter of each peptide as well as the fluorescence of the same diameter directly outside of the area occupied by each peptide. The mean and median fluorescence signal and background pixel intensities as well as other data for each antigen spot were calculated, digitized, and exported as individual rows in a comma-delimited file for subsequent analysis.

### Data processing to identify immunodominant epitopes

A custom script was written to implement a previously-described array processing workflow [32] with a minor change to use the median foreground and background values instead of mean values to minimize outlier effects. Negative background values were interpreted as zeroes. Briefly, background correction was calculated by subtracting the median background from the median foreground measurements for each spot on each array. Normalization was performed by dividing the background-corrected values for each spot on each slide by the non-control spot having the largest fluorescence value on each slide. All spots for each peptide on each array were then summarized into a single value by calculating the median value of the quadruplicate spots for each peptide to further reduce the effects of any outliers. The normalized relative fluorescence intensity values for all peptides and all samples were output as a separate file and how well each peptide was recognized by each of the polyclonal serum samples was quantified.

A separate script was used to transform all relative fluorescence intensity values for each peptide into Z-scores, and separate tables were constructed to contain the summarized Z-score values for all peptides (as columns) representing each of the viral taxa and all samples (as rows) that were tested with the peptide array. A random forest algorithm (randomForest package in R) was applied to each of these tables in order to identify the peptides that were best able to differentiate between each of the viral taxa. In this case, the number of trees generated in the random forest for each species was 100,000, and the number of variables randomly sampled as candidates at each split was equal to the square root of the number of columns present in each table.

The values representing the mean decrease in Gini index were calculated separately for samples obtained from each of the three collections as well as all possible combinations of two or more collections. These data were then used to identify the top 30 peptides according to their usefulness in identifying the correct virus taxon. The BepiPred algorithm was then used to predict the number of residues that are frequently present in B-cell epitopes, and would therefore contribute to increased affinity and binding by antibodies in downstream assays [33]. The peptides were then assigned a cumulative rank based on the epitope prediction and Gini values, and the 10 highest-ranking peptides across the E and NS1 proteins for each viral taxon, as well as 8 peptides in the E2 region for CHIKV, were categorized as the most likely to have high immunodominance and therefore be recognized by antibodies in sera collected from patients around the world. Statistical comparisons of quantitative differences between the Gini and normalized fluorescence values for sets of peptides were performed using a Student’s t-test.

### Human Subject Approval

All samples evaluated on the peptide arrays were acquired from patients under informed consent and approved by the Ethical or Institutional Review Board at each participating institution.

## RESULTS

### Data records

Overall, we screened 137 unique serum samples for their reactivity against a panel of viral peptides (Figure 1, Supplementary Table 1). These samples, together with the clinical diagnosis, were collected from patients with known past exposure to at least one of the viruses targeted by our peptides (Table 1). Text-based data files containing quantified values derived from the raw image data during the peptide array experiments are publicly available in the figshare data repository (https://figshare.com/s/4635efbf387414ce40f7).

**Table 1:** Number of Serum Samples Screened with Peptide Arrays.

| Virus | Number of Samples |
| --- | --- |
| CHIKV | 5 |
| CHIKV, DENV | 32 |
| DENV | 10 |
| DENV1 | 7 |
| DENV2 | 25 |
| DENV3 | 12 |
| WNV | 12 |
| ZIKV | 21 |
| ZIKV, DENV | 9 |
| DENV-Negative | 2 |
| Unknown | 2 |
| Total | 137 |

**Figure 1:**
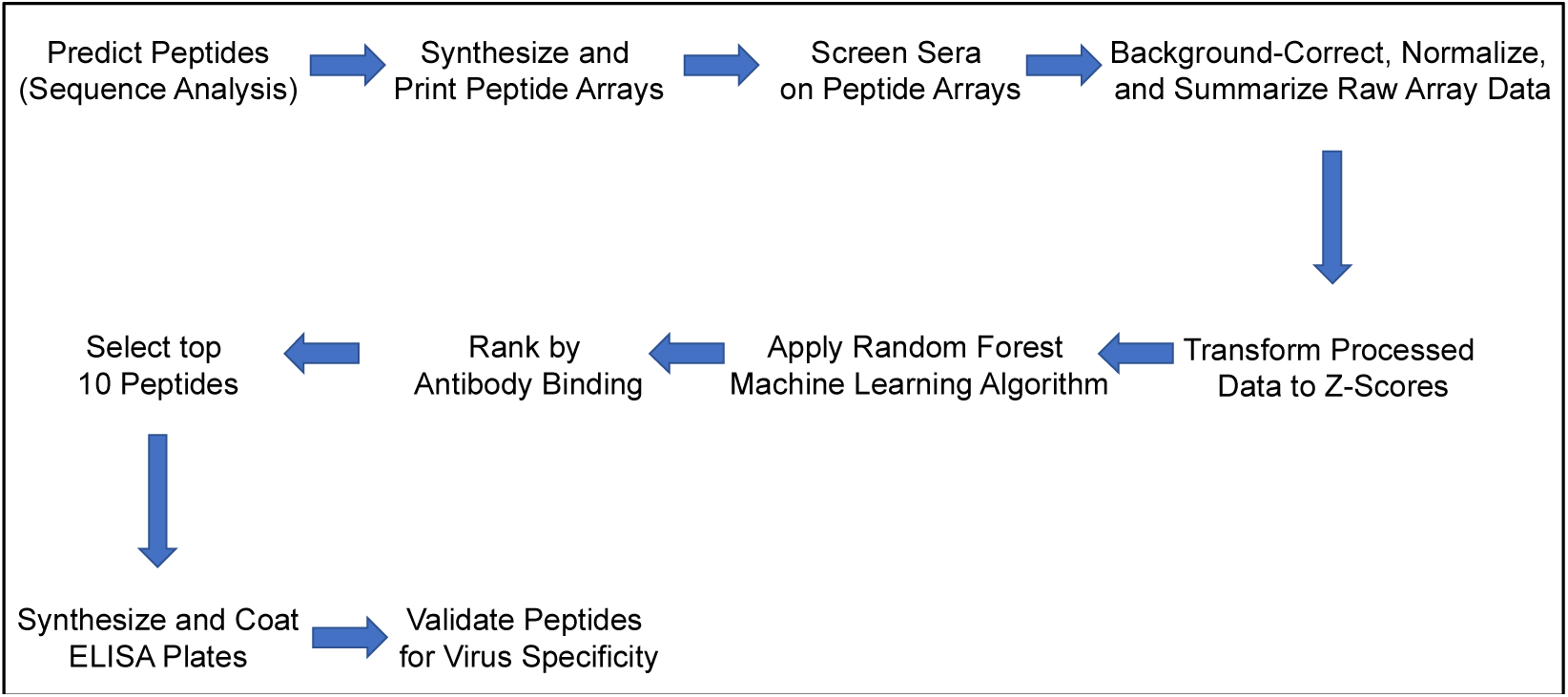
Bioinformatics and Laboratory Workflow Diagram. A graphical depiction of the processes for predicting, screening, processing, and validating peptide array experiments.

The data from each array is contained in a single tab-delimited text file and contains the quantitative data captured from a single serum sample on a single peptide array. A subset of the fields in each file include: location of each peptide spot on the array, peptide identifier, raw mean and median foreground fluorescence at 488 nm, raw mean and median background fluorescence at 488 nm, and other data collected from the raw image.

### Technical Validation

Given the serological cross-reactivity that has been reported among many of our targeted mosquito-borne viruses [34], we recognized the need to validate the results of our high-throughput screen. To do so, we not only ensured that those generating the peptide array data were “blinded” to the phenotype of each sample, but we also computationally evaluated two distinct but complementary quantitative metrics.

First, we compared two serum samples from pediatric patients that had not been infected with DENV prior to sample collection. The data from the DENV-specific peptides in these samples were then compared to those from a representative DENV-positive sample to verify the differences in signal between known positive and known negative samples. This comparison would also provide a better understanding of the contribution of cross-reactivity, which has been reported previously [34], on our platform (Table 2). This comparison showed that the DENV-negative samples had less than four percent of the normalized fluorescence values, well below the 10 percent that was observed in the DENV-positive sample. Transforming these raw data into Z-scores further increases the observed differences in fluorescence values and, provides additional support to the unbiased nature of the data produced in these experiments.

**Table 2:** Comparison of normalized reactivity percentages among representative well-characterized serum samples as an indicator of peptide specificity.

|  | DENV-Negative* |  | DENV-Negative** |  | DENV1-Positive*** |  |
| --- | --- | --- | --- | --- | --- | --- |
|  | DENV | Non-DENV | DENV | Non-DENV | DENV | Non-DENV |
| Min | 0.055% | 0.000% | 0.196% | 0.392% | 0.000% | 0.000% |
| Max | 1.496% | 7.181% | 3.509% | 10.201% | 10.671% | 32.004% |
| Median | 0.383% | 0.550% | 1.032% | 1.854% | 1.455% | 1.576% |
| Mean | 0.505% | 1.005% | 1.242% | 2.186% | 1.837% | 3.576% |

\* Sample H22.
\*\* Sample H23.
\*\*\* Sample H1.

We next wanted to assess the technical rigor of our approach by performing a statistical analysis of the observed experimental variation (Table 3). In this case, data was available for six of our target viruses for which sera was evaluated on the arrays. We specifically wanted to quantify the reactivity of the best-performing peptides for each sample against the target virus. These results show that incorporating Gini scores into our computational pipeline contributed to our ability to identify sets of peptides that were capable of distinguishing between past infection with the majority of our target viruses.

**Table 3:** P-values of comparisons to quantify the difference(s) between the top-performing peptides versus the remaining peptides for each virus taxon to assess experimental variation.

| Virus | Mean Z-score Comparison |  |  |  | Gini Scores of top peptides vs. remaining peptides |
| --- | --- | --- | --- | --- | --- |
|  | Top peptides exposed to positive sera vs. top 10 peptides exposed to negative sera | Top peptides exposed to positive sera vs. remaining peptides exposed to positive sera | Top peptides exposed to positive sera vs. remaining peptides exposed to negative sera | Remaining peptides exposed to positive sera vs. remaining peptides exposed to negative sera |  |
| DENV1 | N.S. | N.S. | N.S. | N.S. | <0.05 |
| DENV2 | N.S. | N.S. | N.S. | N.S. | N.S. |
| DENV3 | <0.05 | N.S. | <0.01 | <0.01 | N.S. |
| CHIKV | <0.01 | <0.05 | <0.001 | <0.001 | <0.001 |
| WNV | N.S. | N.S. | N.S. | <0.05 | <0.05 |
| ZIKV | <0.05 | <0.05 | <0.05 | <0.05 | <0.01 |

It is also important to recognize that each peptide was printed at non-adjacent sites on each array in quadruplicate to minimize experimental bias due to the location of any given spot on the array. Incorporating technical replicates was an important component of the experimental design. Such an approach enables improved replication of the results and also increases the scientific rigor of the resulting dataset upstream of any data processing workflows.

To computationally validate a subset of our high-scoring peptides, we subjected them to a B-cell epitope prediction workflow that incorporates machine learning [35]. Specifically, we selected the best-scoring peptides for each selected taxa and calculated the mean maximum score to be 0.58 (range 0.55 – 0.63). These scores are associated with a specificity greater than 81% (Table 4).

**Table 4:**
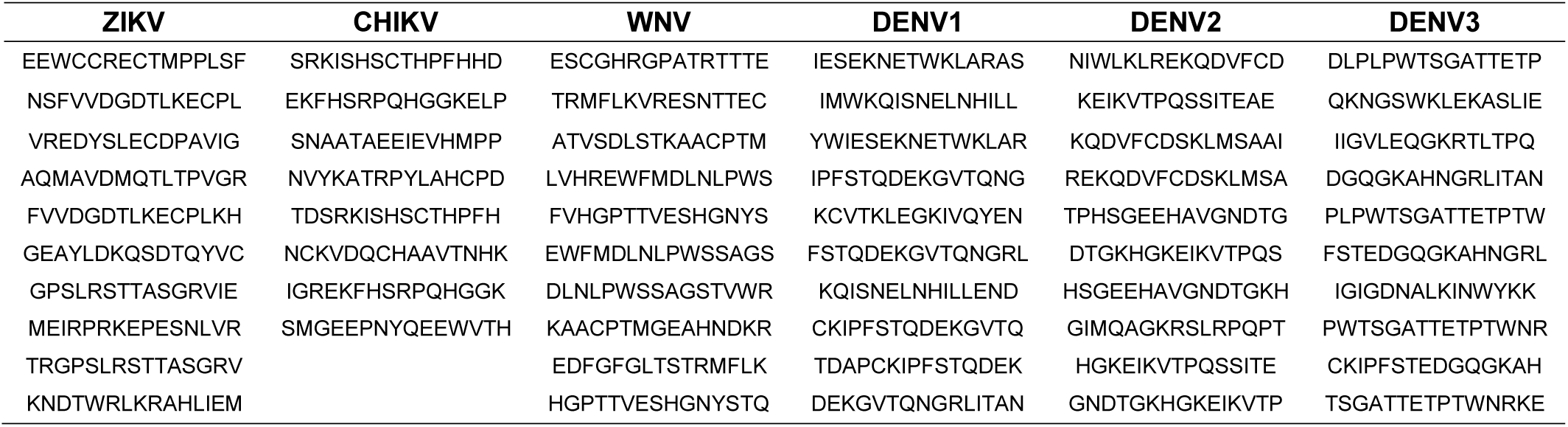
Predicted immunodominant diagnostic epitopes identified from peptide array data reported in this study.

## DISCUSSION

The array data reported in this manuscript were used to identify high-scoring peptides that could be used as serodiagnostic reagents to distinguish between prior infection and seroconversion to a panel of mosquito-borne viruses. Our workflow incorporated both computational and laboratory components to improve identification of regions that were uniquely recognized by virus-specific antibodies to each virus and could therefore be useful as serodiagnostic peptides (Fig. 1). Sabalza *et al.* described a protocol to identify ZIKV specific diagnostic epitopes through peptide microarrays, however, they only used one human serum sample, did not use any bioinformatics analysis, and the identified peptides sequences were not provided [36].

The integration of Gini values calculated by the random forest machine learning algorithm with the BepiPred B-cell epitope prediction algorithm, enabled us to identify the best peptides for each taxon. This approach improved our selected peptides to those that had increased affinity and binding to antibodies [33]. We purposely chose peptides in both the E and NS1 proteins (E2 protein of CHIKV) to improve our ability to detect epitopes within viral antigens that are known to circulate in the bloodstream [11].

We observed that a few of our selected peptides displayed high reactivity and Gini values while other selected peptides had lower measured values. We attribute a subset of these unexpected differences to the imposed requirement of being located within a predicted B-cell epitope. Reactivity is an essential measurement for individual samples, while Gini values are useful to rank peptides based on their ability to identify peptides that differentiate one taxon from the others. As such, Gini values are better able to identify linear epitopes that differentiate taxa and that are sufficiently immunodominant across patient populations. We therefore are quite confident in the results from taxa where the Gini values were significantly different between selected peptides when compared to the remaining peptides.

There were also cases where the comparisons of our selected peptides yielded non-significant p-values when compared to other peptides for the same virus taxon. It is possible that the peptides for these taxa are similarly capable of detecting previous infections, and therefore prevented a significant result. Linear peptides may be unable to adequately differentiate between taxa with non-significant results. A possible alternative is that the peptides are similarly able to differentiate between prior infections, but that the patient sera we tested did not have comprehensive serology for the tested viral pathogens. Given the specificity and sensitivity values that were reported previously for our peptides, we expect this scenario to be more likely. Additional laboratory experiments are being performed to calculate the specificity and sensitivity for our sets of peptides in a larger number of human serum samples.

With these data, it could also be possible to perform the opposite analysis in a way that would search for regions that were recognized with reduced specificity and could therefore be useful to identify peptides that could indicate past infection by at least one of these viruses. Similarly, these data could be mined to identify linear peptides that could be used as antigens to generate an antibody response to such epitopes towards the development of additional “universal” monoclonal antibodies.

These data help to quantify the human humoral response to multiple mosquito-borne viruses and could be useful to identify, map, and/or design native or synthetic antigens that provide increased protection against natural infection by these viruses.

These data could also be relevant to the design of a mosquito-borne virus vaccine. However, care must be taken in designing such experiments to ensure that antibody-dependent enhancement does not increase the risk of adverse events following administering the vaccine.

## ACKNOWLEDGEMENTS

We gratefully acknowledge the Microarray and NGS Core facility at The Scripps Research Institute, especially Shelby Willis, Ryan McBride, Dr. Phillip Ordoukhanian, and Dr. Steven Head for their excellent technical assistance with preparing and developing the peptide arrays. We also thank Dr. Scott Weaver and Dionna Scharton of the World Reference Center for Emerging Viruses and Arboviruses for their assistance with providing control samples.

This project has been funded in whole or part with federal funds from the United States Agency for International Development (USAID) Combating Zika and Future Threats, a Grand Challenge for Development; under grant number AID-OAA-F-16-00102. The Pediatric Dengue Cohort Study from which the Nicaraguan samples were obtained was supported by NIH grants P01 AI106695 (EH), U19AI118610 (EH), R01AI099631 (AB), and Pediatric Dengue Vaccine Initiative grant VE-1 funded by the Bill and Melinda Gates Foundation (EH).

## AUTHOR CONTRIBUTIONS

MPMV contributed to: data analysis and interpretation, manuscript preparation, and editing.

NK and LP Processed and provided human serum samples.

JEC provided human samples from individuals with previous history of natural infection with DENV, ZIKV, or CHIKV or WNV used to generate binding data on the peptide arrays and contributing to manuscript editing. EH, GK, and AB directed the Nicaraguan Pediatric Dengue Cohort Study and provided samples with laboratory-confirmed previous DENV or ZIKV infection; EH contributed to manuscript editing.

BEP contributed to: study conceptualization, experimental design, data analysis and interpretation, manuscript preparation, and editing.

