## Supplemental Table 1 for "Peptide Arrays of Three Collections of Human Sera from Patients Infected with Mosquito-Borne Viruses"

**Supplementary Table 1:** Metadata for Experimental Samples.

| **Sample ID** | **SampleID on Array** | **Protocol 1** | **Protocol 2** | **Protocol 3** | **Data Output File** | **Repository name** |
| --- | --- | --- | --- | --- | --- | --- |
| 527.2B | H1 | Blood Collection | Serum Separation | Peptide Array | JCVI-101_H1_488H50P.txt | figshare |
| 1089.5B | H2 | Blood Collection | Serum Separation | Peptide Array | JCVI-102_H2_488H50P.txt | figshare |
| 5506.7B | H3 | Blood Collection | Serum Separation | Peptide Array | JCVI-103_H3_488H50P.txt | figshare |
| 918.2B | H4 | Blood Collection | Serum Separation | Peptide Array | JCVI-104_H4_488H50P.txt | figshare |
| 5663.7B | H5 | Blood Collection | Serum Separation | Peptide Array | JCVI-51_H5_488H50P.txt | figshare |
| 3383.4B | H6 | Blood Collection | Serum Separation | Peptide Array | JCVI-3_H6_488H50P.txt | figshare |
| 608.8B | H7 | Blood Collection | Serum Separation | Peptide Array | JCVI-4_H7_488H50P.txt | figshare |
| 2347.2B | H8 | Blood Collection | Serum Separation | Peptide Array | JCVI-5_H8_488H50P.txt | figshare |
| 460.5T | H9 | Blood Collection | Serum Separation | Peptide Array | JCVI-6_H9_488H50P.txt | figshare |
| 1264.5B | H10 | Blood Collection | Serum Separation | Peptide Array | JCVI-7_H10_488H50P.txt | figshare |
| 964.3B | H11 | Blood Collection | Serum Separation | Peptide Array | JCVI-8_H11_635H100P_488L50P.txt | figshare |
| 918.3B | H12 | Blood Collection | Serum Separation | Peptide Array | JCVI-95_H12_488H50P.txt | figshare |
| 4191.5B | H13 | Blood Collection | Serum Separation | Peptide Array | JCVI-9_H13_635H100P_488L50P.txt | figshare |
| 1567.3B | H14 | Blood Collection | Serum Separation | Peptide Array | JCVI-10_H14_635H100P_488L50P.txt | figshare |
| 3325.4B | H15 | Blood Collection | Serum Separation | Peptide Array | JCVI-11_H15_635H100P_488L50P.txt | figshare |
| 4394.7B | H16 | Blood Collection | Serum Separation | Peptide Array | JCVI-12_H16_635H100P_488L50P.txt | figshare |
| 1567.2B | H17 | Blood Collection | Serum Separation | Peptide Array | JCVI-13_H17_635H100P_488L50P.txt | figshare |
| 1651.5B | H18 | Blood Collection | Serum Separation | Peptide Array | JCVI-14_H18_635H100P_488L50P.txt | figshare |
| 2898.7B | H19 | Blood Collection | Serum Separation | Peptide Array | JCVI-15_H19_635H100P_488L50P.txt | figshare |
| 3428.7B | H20 | Blood Collection | Serum Separation | Peptide Array | JCVI-16_H20_635H100P_488L50P.txt | figshare |
| 2456.8B | H21 | Blood Collection | Serum Separation | Peptide Array | JCVI-17_H21_635H100P_488L50P.txt | figshare |
| 2088.1B | H22 | Blood Collection | Serum Separation | Peptide Array | JCVI-18_H22_635H100P_488L50P.txt | figshare |
| 3755.1B | H23 | Blood Collection | Serum Separation | Peptide Array | JCVI-19_H23_635H100P_488L50P.txt | figshare |
| 7031.13.A.2 | H24 | Blood Collection | Serum Separation | Peptide Array | JCVI-20_H24_635H100P_488L50P.txt | figshare |
| 4774.13..A.2 | H25 | Blood Collection | Serum Separation | Peptide Array | JCVI-21_H25_635H100P_488L50P.txt | figshare |
| 6681.13.A.2 | H26 | Blood Collection | Serum Separation | Peptide Array | JCVI-22_H26_635H100P_488L50P.txt | figshare |
| 7320.13.A.2 | H27 | Blood Collection | Serum Separation | Peptide Array | JCVI-23_H27_635H100P_488L50P.txt | figshare |
| 4723.13.A.2 | H28 | Blood Collection | Serum Separation | Peptide Array | JCVI-24_H28_635H100P_488L50P.txt | figshare |
| 6215.13.A.2 | H29 | Blood Collection | Serum Separation | Peptide Array | JCVI-25_H29_635H100P_488L50P.txt | figshare |
| 4858.13.A.2 | H30 | Blood Collection | Serum Separation | Peptide Array | JCVI-26_H30_635H100P_488L50P.txt | figshare |
| 5604.13.A.2 | H31 | Blood Collection | Serum Separation | Peptide Array | JCVI-28_H31_635H100P_488L50P.txt | figshare |
| 7420.13.A.2 | H32 | Blood Collection | Serum Separation | Peptide Array | JCVI-27_H32_635H100P_488L50P.txt | figshare |
| D1 | L1 | Blood Collection | Serum Separation | Peptide Array | JCVI-29_L1_635H100P_488L50P.txt | figshare |
| D2 | L2 | Blood Collection | Serum Separation | Peptide Array | JCVI-30_L2_635H100P_488L50P.txt | figshare |
| D3 | L3 | Blood Collection | Serum Separation | Peptide Array | JCVI-31_L3_635H100P_488L50P.txt | figshare |
| D4 | L4 | Blood Collection | Serum Separation | Peptide Array | JCVI-32_L4_635H100P_488L50P.txt | figshare |
| D5 | L5 | Blood Collection | Serum Separation | Peptide Array | JCVI-33_L5_635H100P_488L50P.txt | figshare |
| D6 | L6 | Blood Collection | Serum Separation | Peptide Array | JCVI-34_L6_635H100P_488L50P.txt | figshare |
| D7 | L7 | Blood Collection | Serum Separation | Peptide Array | JCVI-35_L7_635H100P_488L50P.txt | figshare |
| D8 | L8 | Blood Collection | Serum Separation | Peptide Array | JCVI-36_L8_635H100P_488L50P.txt | figshare |
| D9 | L9 | Blood Collection | Serum Separation | Peptide Array | JCVI-37_L9_635H100P_488L50P.txt | figshare |
| D10 | L10 | Blood Collection | Serum Separation | Peptide Array | JCVI-38_L10_635H100P_488L50P.txt | figshare |
| D11 | L11 | Blood Collection | Serum Separation | Peptide Array | JCVI-39_L11_635H100P_488L50P.txt | figshare |
| D12 | L12 | Blood Collection | Serum Separation | Peptide Array | JCVI-40_L12_635H100P_488L50P.txt | figshare |
| D13 | L13 | Blood Collection | Serum Separation | Peptide Array | JCVI-41_L13_635H100P_488L50P.txt | figshare |
| D14 | L14 | Blood Collection | Serum Separation | Peptide Array | JCVI-42_L14_635H100P_488L50P.txt | figshare |
| D15 | L15 | Blood Collection | Serum Separation | Peptide Array | JCVI-43_L15_635H50P_488L50P.txt | figshare |
| D16 | L16 | Blood Collection | Serum Separation | Peptide Array | JCVI-44_L16_635H100P_488L50P.txt | figshare |
| D17 | L17 | Blood Collection | Serum Separation | Peptide Array | JCVI-45_L17_635H100P_488L50P.txt | figshare |
| D18 | L18 | Blood Collection | Serum Separation | Peptide Array | JCVI-46_L18_635H100P_488L50P.txt | figshare |
| D19 | L19 | Blood Collection | Serum Separation | Peptide Array | JCVI-47_L19_635H100P_488L50P.txt | figshare |
| D20 | L20 | Blood Collection | Serum Separation | Peptide Array | JCVI-48_L20_635H100P_488L50P.txt | figshare |
| D21 | L21 | Blood Collection | Serum Separation | Peptide Array | JCVI-49_L21_635H100P_488L50P.txt | figshare |
| D22 | L22 | Blood Collection | Serum Separation | Peptide Array | JCVI-50_L22_635H100P_488L50P.txt | figshare |
| D23 | L23 | Blood Collection | Serum Separation | Peptide Array | JCVI-51_L23_635H100P_488L50P.txt | figshare |
| Z1 | L24 | Blood Collection | Serum Separation | Peptide Array | JCVI-52_L24_635H100P_488L50P.txt | figshare |
| Z2 | L25 | Blood Collection | Serum Separation | Peptide Array | JCVI-54_L25_635H100P_488L50P.txt | figshare |
| Z3 | L26 | Blood Collection | Serum Separation | Peptide Array | JCVI-55_L26_635H100P_488L50P.txt | figshare |
| Z4 | L27 | Blood Collection | Serum Separation | Peptide Array | JCVI-56_L27_635H100P_488L50P.txt | figshare |
| Z5 | L28 | Blood Collection | Serum Separation | Peptide Array | JCVI-57_L28_635H100P_488L50P.txt | figshare |
| Z6 | L29 | Blood Collection | Serum Separation | Peptide Array | JCVI-58_L29_635H75P_488L50P.txt | figshare |
| Z7 | L30 | Blood Collection | Serum Separation | Peptide Array | JCVI-59_L30_635H100P_488L50P.txt | figshare |
| Z8 | L31 | Blood Collection | Serum Separation | Peptide Array | JCVI-60_L31_635H100P_488L50P.txt | figshare |
| Z9 | L32 | Blood Collection | Serum Separation | Peptide Array | JCVI-61_L32_635H100P_488L50P.txt | figshare |
| Donor 976 | C1 | Blood Collection | Serum Separation | Peptide Array | JCVI-99_C1_488H50P.txt | figshare |
| Donor 977 | C2 | Blood Collection | Serum Separation | Peptide Array | JCVI-62_C2_635H100P_488L50P.txt | figshare |
| Donor 978 | C3 | Blood Collection | Serum Separation | Peptide Array | JCVI-63_C3_635H100P_488L50P.txt | figshare |
| Donor 878 | C4 | Blood Collection | Serum Separation | Peptide Array | JCVI-65_C4_635H100P_488L50P.txt | figshare |
| Donor 879 | C5 | Blood Collection | Serum Separation | Peptide Array | JCVI-66_C5_635H100P_488L50P.txt | figshare |
| Donor 880 | C6 | Blood Collection | Serum Separation | Peptide Array | JCVI-67_C6_635H100P_488L50P.txt | figshare |
| Donor 881 | C7 | Blood Collection | Serum Separation | Peptide Array | JCVI-68_C7_635H100P_488L50P.txt | figshare |
| Donor 882 | C8 | Blood Collection | Serum Separation | Peptide Array | JCVI-69_C8_635H100P_488L50P.txt | figshare |
| Donor 883 | C9 | Blood Collection | Serum Separation | Peptide Array | JCVI-71_C9_635H100P_488L50P.txt | figshare |
| Donor 884 | C10 | Blood Collection | Serum Separation | Peptide Array | JCVI-72_C10_635H100P_488L50P.txt | figshare |
| Donor 885 | C11 | Blood Collection | Serum Separation | Peptide Array | JCVI-73_C11_635H100P_488L50P.txt | figshare |
| Donor 886 | C12 | Blood Collection | Serum Separation | Peptide Array | JCVI-74_C12_635H100P_488L50P.txt | figshare |
| Donor 887 | C13 | Blood Collection | Serum Separation | Peptide Array | JCVI-75_C13_635H100P_488L50P.txt | figshare |
| Donor 888 | C14 | Blood Collection | Serum Separation | Peptide Array | JCVI-76_C14_635H100P_488L50P.txt | figshare |
| Donor 889 | C15 | Blood Collection | Serum Separation | Peptide Array | JCVI-77_C15_635H100P_488L50P.txt | figshare |
| Donor 890 | C16 | Blood Collection | Serum Separation | Peptide Array | JCVI-78_C16_635H100P_488L50P.txt | figshare |
| Donor 891 | C17 | Blood Collection | Serum Separation | Peptide Array | JCVI-79_C17_635H100P_488L50P.txt | figshare |
| Donor 892 | C18 | Blood Collection | Serum Separation | Peptide Array | JCVI-80_C18_635H100P_488L50P.txt | figshare |
| Donor 893 | C19 | Blood Collection | Serum Separation | Peptide Array | JCVI-81_C19_635H50P_488L50P.txt | figshare |
| Donor 894 | C20 | Blood Collection | Serum Separation | Peptide Array | JCVI-82_C20_635H100P_488L50P.txt | figshare |
| Donor 895 | C21 | Blood Collection | Serum Separation | Peptide Array | JCVI-83_C21_635H100P_488L50P.txt | figshare |
| Donor 896 | C22 | Blood Collection | Serum Separation | Peptide Array | JCVI-84_C22_635H100P_488L50P.txt | figshare |
| Donor 897 | C23 | Blood Collection | Serum Separation | Peptide Array | JCVI-85_C23_635H100P_488L50P.txt | figshare |
| Donor 898 | C24 | Blood Collection | Serum Separation | Peptide Array | JCVI-97_C24_635H50P_488L50P.txt | figshare |
| Donor 899 | C25 | Blood Collection | Serum Separation | Peptide Array | JCVI-98_C25_635H50P_488L50P.txt | figshare |
| Donor 900 | C26 | Blood Collection | Serum Separation | Peptide Array | JCVI-99_C26_635H50P_488L50P.txt | figshare |
| Donor 901 | C27 | Blood Collection | Serum Separation | Peptide Array | JCVI-100_C27_635H50P_488L50P.txt | figshare |
| Donor 902 | C28 | Blood Collection | Serum Separation | Peptide Array | JCVI-101_C28_635H50P_488L50P.txt | figshare |
| Donor 938 | C29 | Blood Collection | Serum Separation | Peptide Array | JCVI-104_C29_635H50P_488L50P.txt | figshare |
| Donor 939 | C30 | Blood Collection | Serum Separation | Peptide Array | JCVI-106_C30_635H50P_488L50P.txt | figshare |
| Donor 947 | C31 | Blood Collection | Serum Separation | Peptide Array | JCVI-114_C31_635H50P_488L50P.txt | figshare |
| Donor 948 | C32 | Blood Collection | Serum Separation | Peptide Array | JCVI-115_C32_635H50P_488L50P.txt | figshare |
| Donor 949 | C33 | Blood Collection | Serum Separation | Peptide Array | JCVI-116_C33_635H50P_488L50P.txt | figshare |
| Donor 950 | C34 | Blood Collection | Serum Separation | Peptide Array | JCVI-117_C34_635H50P_488L50P.txt | figshare |
| Donor 951 | C35 | Blood Collection | Serum Separation | Peptide Array | JCVI-131_C35_635H50P_488L50P.txt | figshare |
| Donor 952 | C36 | Blood Collection | Serum Separation | Peptide Array | JCVI-138_C36_635H50P_488L50P.txt | figshare |
| Donor 953 | C37 | Blood Collection | Serum Separation | Peptide Array | JCVI-139_C37_635H50P_488L50P.txt | figshare |
| Donor 954 | C38 | Blood Collection | Serum Separation | Peptide Array | JCVI-140_C38_635H50P_488L50P.txt | figshare |
| Donor 955 | C39 | Blood Collection | Serum Separation | Peptide Array | JCVI-142_C39_635H50P_488L50P.txt | figshare |
| Donor 956 | C40 | Blood Collection | Serum Separation | Peptide Array | JCVI-102_C40_635H50P_488L50P.txt | figshare |
| Donor 160 | C41 | Blood Collection | Serum Separation | Peptide Array | JCVI-98_C41_488L50P.txt | figshare |
| Donor 243 | C42 | Blood Collection | Serum Separation | Peptide Array | JCVI-108_C42_635H50P_488L50P.txt | figshare |
| Donor 244 | C43 | Blood Collection | Serum Separation | Peptide Array | JCVI-109_C43_635H50P_488L50P.txt | figshare |
| Donor 338 | C44 | Blood Collection | Serum Separation | Peptide Array | JCVI-111_C44_635H50P_488L50P.txt | figshare |
| Donor 538 | C45 | Blood Collection | Serum Separation | Peptide Array | JCVI-112_C45_635H50P_488L50P.txt | figshare |
| Donor 848 | C48 | Blood Collection | Serum Separation | Peptide Array | JCVI-124_C48_635H50P_488L50P.txt | figshare |
| Donor 849 | C49 | Blood Collection | Serum Separation | Peptide Array | JCVI-155_C49_635H50P_488L50P.txt | figshare |
| Donor 960 | C50 | Blood Collection | Serum Separation | Peptide Array | JCVI-158_C50_635H50P_488L50P.txt | figshare |
| Donor 865 | C52 | Blood Collection | Serum Separation | Peptide Array | JCVI-130_C52_635H50P_488L50P.txt | figshare |
| Donor 866 | C53 | Blood Collection | Serum Separation | Peptide Array | JCVI-132_C53_635H50P_488L50P.txt | figshare |
| Donor 867 | C54 | Blood Collection | Serum Separation | Peptide Array | JCVI-96_C54_635H50P_488L50P.txt | figshare |
| Donor 868 | C55 | Blood Collection | Serum Separation | Peptide Array | JCVI-86_C55_635H50P_488L50P.txt | figshare |
| Donor 869 | C56 | Blood Collection | Serum Separation | Peptide Array | JCVI-87_C56_635H50P_488L50P.txt | figshare |
| Donor 870 | C57 | Blood Collection | Serum Separation | Peptide Array | JCVI-88_C57_635H50P_488L50P.txt | figshare |
| Donor 871 | C58 | Blood Collection | Serum Separation | Peptide Array | JCVI-90_C58_635H50P_488L50P.txt | figshare |
| Donor 872 | C59 | Blood Collection | Serum Separation | Peptide Array | JCVI-91_C59_635H50P_488L50P.txt | figshare |
| Donor 873 | C60 | Blood Collection | Serum Separation | Peptide Array | JCVI-93_C60_635H50P_488L50P.txt | figshare |
| Donor 874 | C61 | Blood Collection | Serum Separation | Peptide Array | JCVI-94_C61_635H50P_488L50P.txt | figshare |
| Donor 875 | C62 | Blood Collection | Serum Separation | Peptide Array | JCVI-95_C62_635H50P_488L50P.txt | figshare |
| Donor 876 | C63 | Blood Collection | Serum Separation | Peptide Array | JCVI-102_C63_635H50P_488L50P.txt | figshare |
| Donor 1002 | C64 | Blood Collection | Serum Separation | Peptide Array | JCVI-110_C64_635H50P_488L50P.txt | figshare |
| Donor 1008 | C65 | Blood Collection | Serum Separation | Peptide Array | JCVI-113_C65_635H50P_488L50P.txt | figshare |
| Donor 1010 | C66 | Blood Collection | Serum Separation | Peptide Array | JCVI-118_C66_635H50P_488L50P.txt | figshare |
| Donor 1011 | C67 | Blood Collection | Serum Separation | Peptide Array | JCVI-119_C67_635H50P_488L50P.txt | figshare |
| Donor 1012 | C68 | Blood Collection | Serum Separation | Peptide Array | JCVI-150_C68_635H50P_488L50P.txt | figshare |
| Donor 1016 | C69 | Blood Collection | Serum Separation | Peptide Array | JCVI-135_C69_635H50P_488L50P.txt | figshare |
| Donor 1046 | C70 | Blood Collection | Serum Separation | Peptide Array | JCVI-100_C70_488H50P.txt | figshare |
| Donor 1057 | C71 | Blood Collection | Serum Separation | Peptide Array | JCVI-137_C71_635H50P_488L50P.txt | figshare |
| Donor 1060 | C72 | Blood Collection | Serum Separation | Peptide Array | JCVI-153_C72_635H50P_488L50P.txt | figshare |
| Donor 1062 | C73 | Blood Collection | Serum Separation | Peptide Array | JCVI-152_C73_635H50P_488L50P.txt | figshare |
| Donor 1063 | C74 | Blood Collection | Serum Separation | Peptide Array | JCVI-151_C74_635H50P_488L50P.txt | figshare |
| Donor 1064 | C75 | Blood Collection | Serum Separation | Peptide Array | JCVI-133_C75_635H50P_488L50P.txt | figshare |
| Donor 973 | C76 | Blood Collection | Serum Separation | Peptide Array | JCVI-157_C76_635H50P_488L50P.txt | figshare |
